# KRAS^G12D^ and TP53^R167H^ Cooperate to Induce Pancreatic Ductal Adenocarcinoma in *Sus Scrofa* Pigs

**DOI:** 10.1101/358416

**Authors:** Daniel R. Principe, Nana Haahr Overgaard, Alex J. Park, Andrew M. Diaz, Carolina Torres, Ronald McKinney, Matthew J. Dorman, Karla Castellanos, Regina Schwind, David W. Dawson, Ajay Rana, Ajay Maker, Hidayatullah G. Munshi, Laurie A. Rund, Paul J. Grippo, Lawrence B. Schook

## Abstract

Although survival has improved in recent years, the prognosis of patients with advanced pancreatic ductal adenocarcinoma (PDAC) remains poor. Despite substantial differences in anatomy, physiology, genetics, and metabolism, the overwhelming majority of preclinical testing relies on transgenic mice. Hence, while mice have allowed for tremendous advances in cancer biology, they have been a poor predictor of drug performance/toxicity in the clinic. Given the greater similarity of *sus scrofa* pigs to humans, we engineered transgenic *sus scrofa* expressing a LSL-KRAS^G12D^-TP53^R167H^ cassette. By applying Adeno-Cre to pancreatic duct cells *in vitro*, cells self-immortalized and established tumors in immunocompromised mice. When Adeno-Cre was administered to the main pancreatic duct *in vivo*, pigs developed extensive PDAC at the injection site hallmarked by excessive proliferation and desmoplastic stroma. This serves as the first large animal model of pancreatic carcinogenesis, and may allow for insight into new avenues of translational research not before possible in rodents.

## Introduction

Genetically modified mouse models have revolutionized cancer research, allowing for faithful and reproducible histotypes closely representing those observed in human patients. This is particularly true in recent years, largely attributed to the availability of mixand-match systems such as Cre/lox and tTA/TRE^1^. These respective technologies continue to provide tissue specific recombination and gene inducibility, while allowing for more clinically relevant models of non-syndromic cancers. By employing such model systems, mice have offered invaluable and unprecedented insight into the etiology of diseases that generally present in late stages, such as pancreatic ductal adenocarcinoma (PDAC)^2^. However, while mice have inarguably allowed for tremendous advances in cancer, they have considerable limitations as a disease model due to vastly different physiology^3^, and the conceptual leap from mouse to human has been a culprit in the failure of many promising preclinical therapies.

The most obvious of these limitations is the mouse anatomy. The average human is roughly 2,500-3,000 times the size of an average mouse, severely limiting the use of mice for studies focused on surgical or radiologic interventions. Importantly, the domestic pig (*sus scrofa*) is now emerging as a viable model for cancer and many other human diseases. While the mouse (*Mus musculus*) genome is approximately 14% smaller than that of humans (2.5Gb compared to 2.9Gb)^4^, the *sus scrofa* genome is more similar at 2.7Gb^5,6^ Furthermore, the linkage conservation between humans and pigs is considerably more extensive than that between humans and mice^7^. As a result, pigs are more closely related to humans with respect to anatomy, physiology, and immunology^8^, and have the potential to more accurately represent a variety of human diseases, particularly those affecting the pancreas.

The porcine pancreas is far more human-like than that of rodents, with comparable anatomical orientation and localization^9^. For instance, pig pancreata generally exhibit three distinct lobes, with the duodenal lobe corresponding to the head of the human pancreas, the connecting lobe being analogous to the uncinate process, and the splenic lobe corresponding to the tail/body of the pancreas^9^. In light of these many similarities, modeling pancreatic cancer in pigs would provide the opportunity to conduct surgical and radiologic studies never before possible, and may offer additional insight into disease pathogenesis due to the size and composition of the gland.

Currently, the most widely used mouse model of pancreatic cancer consists of concurrent KRAS^G12D^ and TP53172H mutations targeted to the exocrine pancreas under the Pdx-1 promoter^10^. This model (KPC) has been used extensively and is considered a gold standard within the field. We sought to generate a similar histoype in *sus scrofa* pigs to provide a more physiologically and anatomically relevant model. We first generated pigs harboring analogous KRAS^G12D^ and TP53R167H mutations, which are observed in approximately 95% and 50% of human PDAC patients respectively^11,12^. After demonstrating the ability of these transgenes to induce adenocarcinomatous transformation of pig duct cells *ex vivo*, we administered an Adenoviral Cre Recombinase (Adeno-Cre) to the pig pancreas *in vivo*. Following refinement of the delivery system, these mutations successfully induced pancreatic neoplasms and cancer in the pig, providing the first documented large animal model of pancreatic carcinoma.

## Materials and Methods

### Cloning and sequencing of porcine KRAS and TP53 genes in TOPO shuttle vector

Porcine bone marrow cells were isolated and frozen at −80°C. Total RNA was extracted from these cells with RNeasy mini kit (QIAGEN, Hilden, Germany) and reverse transcribed into cDNA with the QuantiTect Reverse Transcription Kit (QIAGEN). Ensembl genome browser was used to design the PCR primer sequences for amplification of porcine-specific KRAS (ENSSSCG00000000561) and TP53 genes (ENSSSCT00000019534). The forward and reverse primer sequences for KRAS were 5’-CTGCTGAAAATGACTGAATATAAACTT-3’ and 5’-TTACATAATTATACACTTTGTCTTTGA-3’, respectively. The PCR thermal cycling conditions for KRAS amplification were 94°C for 10 min, followed by 30 cycles of 94°C for 30 sec, 55°C for 3 min, and 65°C for 1 min with a final 72°C step for 10 min. The forward and reverse primer sequences used for TP53 were 5’-TGCAATGGCGGAGTCGCAG-3’ and 5’-TCAGTCTGAGTCAGGTCCTTC-3’, respectively.

The PCR thermal cycling conditions were 94°C for 10 min, followed by 30 cycles of 94°C for 30 sec, 55°C for 30 sec, and 68°C for 1 min with a final 72°C step for 10 min. ACTB was used as an endogenous control. The forward and reverse primer sequences used were 5’-GACATCCGCAAGGACCTCTA-3’ and 5’-ACATCTGCTGGAAGGTGGAC-3’, respectively. The PCR conditions used were the same as for TP53. PCR amplified KRAS and TP53 cDNAs were then cloned into pCR2.1-TOPO vectors using the TOPO TA cloning kit according to the manufacturer’s instructions (Invitrogen, Waltham, MA) and the cDNAs confirmed to have 100% nucleotide identity to porcine KRAS and TP53 (http://useast.ensembl.org/index.html).

### Site-directed mutagenesis of KRAS and TP53 cDNA

QuikChange Site-Directed Mutagenesis Kit (Stratagene, La Jolla, CA) was used to introduce changes to the nucleotide sequence corresponding to the G12D mutation into the cloned porcine KRAS cDNA and the R167H mutation into the cloned porcine TP53 cDNA. The primers used to generate the G12D mutation in KRAS were 5’-TGGTAGTTGGAGCTGATGGCGTAGGCAAGAG-3’ and 5’-CTCTTGCCTACGCCATCAGCTCCAACTACCA-3’. The primers used to generate the R167H mutation in TP53 were 5’-GAGGTGGTGAGGCACTGTCCCCACCAT-3’ and 5’-ATGGTGGGGACAGTGCCTCACCACCTC-3’. Mutations were confirmed by sequencing.

### Cloning of KRAS^G12D^ and TP53^R167H^ cDNAs into the pIRES vector

The aforementioned KRASG12D cDNA was PCR amplified with primers 5’-CTAGCTAGCTAGCTGCTGAAAATGACTGAATAT-3’ and 5’-CCGCTCGAGCGGTTACATAATTATACAC-3’ for 30 cycles of 94°C for 1 min, 94°C for 30 sec, 60°C for 1 min, and 68°C for 1 min, and the resultant fragment cloned into the NheI and XhoI sites of pIRES (Clontech, CA). The aforementioned TP53R167H cDNA was PCR amplified with primers 5’-ACGCGTGGACGTCTTGGCCATATGCAATGGAGGA3’ and 5’-ATAAGAATGCGGCCGCTAAACTATTCAGTCTGAGTCAGGTCC-3’ for 30 cycles of 94°C for 1 min, 94°C for 30 sec, 65°C for 1 min, and 68°C for 1 min, and the resultant fragment cloned into the SalI and NotI sites of pIRES containing the KRASG12D cDNA. cDNA sequences of KRAS^G12D^ and TP53^R167H^ were confirmed correct in the resultant vector by sequencing.

### Cloning CAG promoter into a loxP-STOP(polyA)-loxP containing vector

The pkw15 plasmid containing the CAG promoter and the pkw13 plasmid containing LoxP-STOP-(polyA)-loxP were used for the vector construction. A CAG promoter was isolated by SpeI/MfeI digestion and cloned into the pkw13 plasmid at SpeI/EcoRI cloning sites.

### Construction of the inducible KRAS^G12D^ and TP53^R167H^ oncopig expression vector

The KRAS^G12D^-TP53^R167H^-pIRES vector was digested with PvuI and NheI, and the resulting KRAS^G12D^-IRES-TP53^R167H^-polyA fragments were gel purified and cloned into CAG-LSL-pkw13 vectors at PacI/NheI sites. cDNA sequences of KRAS^G12D^ and TP53^R167H^ in the final vector were confirmed correct in the resultant vector by sequencing.

### Generation of fetal fibroblast strains

Male fetal fibroblasts cells (FFCs) from Minnesota miniature pigs (NSRRC:0005) were collected as described^13^ with minor modification. Briefly, after removing the head and internal organs, the fetus was minced and digested individually in 20 ml of digestion media (Dulbecco’s Modified Eagle Medium (DMEM) supplemented with 15% fetal bovine serum (FBS), 200 units/ml collagenase and 25 units/ml DNaseI) for 4–5 hours at 38.5°C and 5% CO_2_ in air. Digested cells were washed with DMEM supplemented with 15% FBS (Hyclone, Logan, UT) and 10μg/ml gentamicin, cultured overnight, and then collected and frozen at −80°C in FBS supplemented with 10% dimethyl sulfoxide (DMSO) (v/v) and stored in liquid nitrogen.

### Production of transgenic cells

Early passage number FFCs (P1-2) were cultured in cell culture medium (DMEM supplemented with 15% (v/v) FBS, 2.5 ng/ml basic fibroblast growth factor and 10μg/ml gentamicin) overnight and grown to 75–85% confluency. Media was replaced 4 hours prior to transfection. FFCs were washed for 1–2 min with phosphate buffered saline (PBS; Invitrogen) and harvested with 0.05% trypsin-EDTA (Invitrogen; 1 ml per 75cm^2^ flask). Cells were resuspended in cell culture medium, pelleted at 600 × g for 10 min, resuspended in 10 ml Opti-MEM (Invitrogen), and then quantified using a hemocytometer and re-pelleted. Cells were resuspended in transfection media (75% cytosalts [120mMKCl, 0.15mM CaCl_2_, 10mM K2HPO4; pH 7.6, 5mM MgCl_2_]^14^ and 25% Opti-MEM [Gibco BRL, Grand Island, NY]). The cell concentration was adjusted to 1×106 cells/ml and 200μl of cells were co-transfected by electroporation with linearized mutant *KRAS* and *TP53* construct containing a Neo selectable casette (2μg). Electroporation utilized three consecutive 250-V, 1-ms square wave pulses administered through a BTX ECM 2001 (BTX, San Diego, CA) in a 2mM gap cuvette. After electroporation, cells were plated in a 100mM dish at 3,000 cells per dish in cell culture medium. After 36 hours, cells were selected by the addition of geneticin (G418; 400μg/ml) for 10–14 days until the formation of cell colonies. Genomic DNA from the cell colonies was used to verify the presence of both transgenes by PCR. These cells then were stored in liquid nitrogen until used as donor cells for SCNT.

### Oocyte maturation, SCNT, and embryo reconstruction

Fibroblast cells identified to have integration of the transgenes (*KRAS* and *TP53*) were used as donor cells for SCNT into enucleated oocytes followed by electrical fusion and activation as previously described^15^. In brief, cumulus-oocyte cell complexes (COCs) were received in Phase I maturation medium from ART Inc. (Madison, WI) approximately 24 hours after harvest. COCs were then cultured in fresh Phase II maturation medium for a total of 40 hours in a humidified atmosphere of 5% CO_2_ at 38.5°C. Phase I and II medium were supplied by ART Inc. Expanded COCs were then vortexed in 0.1% hyaluronidase in Hepes-buffered Tyrode’s medium containing 0.01% PVA for 4 min to remove the cumulus cells. Oocytes having a visible first polar body (PB) with uniform cytoplasm were selected and placed in fresh manipulation medium (25mM Hepes-buffered TCM199 with 3 mg/ml BSA) containing 7.5μg/ml cytochalasin B that was overlaid with warm mineral oil.

The PB, MII chromosomes, and a small amount of surrounding cytoplasm of the oocyte were enucleated using a beveled glass pipette with an inner diameter of 17–20 μm. After enucleation, a donor cell was injected into the perivitelline space and placed adjacent to the recipient cytoplasm. Karyoplast–cytoplast complexes were fused and activated with 2 DC pulses (1 sec interval) of 1.2 kV/cm for 30 μsec provided by a BTX Electro-cell Manipulator 200 in fusion medium (0.3 M mannitol, 1.0mM CaCl2, 0.1mM MgCl2, and 0.5mM Hepes, pH adjusted to 7.0–7.4). After simultaneous fusion and activation, only the fused embryos were cultured into four well cell plates (Nunc, Denmark) containing 500μl of PZM3 with 0.3% BSA and 500nM Scriptaid at 38.5°C and 5% CO2 in humidified air for 14 to 16 hours, until embryo transfer.

### Embryo transfer and piglet production

More than 100 SCNT zygotes were surgically transferred to the oviducts of surrogates on the day of, or one day after, the onset of estrus. The pregnant surrogates were monitored via ultrasound throughout pregnancy. Piglets were delivered via cesarean section from surrogates by day 114–116 of gestation. Piglets are processed immediately and tissue samples were collected for establishment of cell lines and PCR genotyping. Piglets were then hand-raised until weaning (3–4 weeks of age).

### Pigs and In Vivo Imaging

All animal procedures were approved by the University of Illinois Institutional Animal Care and Use Committee. All pigs used were of the LSL-KRAS^G12D^-TP53^R167H^ transgenic line as described previously^16^. LSL-KRAS^G12D^-TP53^R167H^ pigs were dosed with 4×109 PFU of Ad5CMVCre-eGFP (Adeno-Cre), either directly into the body of the pancreas or into the pancreatic duct pushed toward the pancreas. For all cases the pancreas was accessed surgically through a ventral midline incision. Anesthesia was induced with TKX (Telazol, Ketamine, Xylazine), intubated and maintained in a surgical plane of anesthesia with Isoflurane.

Following recovery, all animals were monitored for general health, weight and periodic blood chemistry evaluation. For euthanasia, each animal was sedated and euthanized with sodium pentobarbital overdose (Fatal plus 10 cc/100 lbs.). All animals underwent gross and histopathologic evaluation. Tissues were then collected and subject to gross and histopathologic analysis.

For *in vivo* imaging, animals were sedated as described previously and 120ml of the contrast agent Iohexal (Omnipaque) was delivered intravenously, allowing for visualization of potential pathologies by contrast computed tomography (CT). Scans were limited to the cranial abdomen.

### Cell Culture

The pancreas from a two-month old female pig was collected and immediately cut into 1mm^3^ pieces in ice cold DMEM/F12 media with 20% animal serum complex, penicillin (100 units/mL), and streptomycin (100 [.proportional]g/mL). Tissue was transferred to 50ml conical tubes and washed with cold media and transferred to a sterile hood. Tissues were washed again and incubated with Collagenase Ia (1 mg/ml) and 0.25 mg/ml of trypsin inhibitor for five minutes at 37 °C. The Collagenase reaction was quenched with PBS with 20% animal serum complex and the cells washed. The reaction was similarly quenched with PBS with 20% animal serum complex and the cells washed in fresh culture media with 20% heat-inactivated fetal bovine serum (FBS), penicillin (100 units/mL), and streptomycin (100 [.proportional]g/mL). The cells were then seeded on six well plates and allowed to adhere. Once the cells were seeded, they were infected with Adeno-Cre at a 200 to 500 MOI. The media was changed after six hours and cells maintained at 37 °C with 5% CO2.

### Mice

5×10^6^ PORC1 cells suspended in100μl of Matrigel, (BD Biosciences, San Diego, CA, USA) and injected subcutaneously into 3 SCID mice (NOD.CB17-Prkdcscid/ JAX, Bar Harbor, ME, USA). Tumor growth data was monitored for up to 100 days. SQ tumors were routinely measured and overall health was assessed by weight and visual inspection. Mice were sacrificed by cervical dislocation, and tissues collected and subject to histopathology. All animal procedures were approved by the University of Illinois Institutional Animal Care and Use Committee.

### RT-PCR

RNA was extracted using Trizol (Fischer-Scientific, Waltham, MA). Quantity/quality was evaluated by spectrophotometer (Nanogen Inc, San Diego, CA) and integrity confirmed via gel electrophoresis. cDNA was then synthesized using the High-Capacity cDNA Reverse Transcription Kit per manufacturer specifications (Fischer-Scientific, Waltham, MA) and rtPCR was conducted with SYBR Green and commercially available TP53^R167H^ and GAPDH primers (Fischer-Scientific, Waltham, MA). CAG-F: 5’-TCATATGCCAAGTACGCCCC-3’; CAG-R: 5’-CCCCATCGCTGCACAAAATA-3’; TP53-F: 5’-TGGCTCTCCTCAAGCGTATT-3’; TP53-R: 5’-ATTTTCATCCAGCCAGTTCG-3’. Bands displayed were run as shown and without modification.

### Histology, immunohistochemistry, and immunofluorescence/immunocytochemistry

LSL-KRAS^G12D^-TP53^R167H^ pigs or were euthanized and subjected to pathological examination the pancreas, small intestine, liver, bile duct, lymph nodes, spleen, omenteum, colon, thyroid, lung, heart, kindeys, diaphragm, esophagus, bladder, and stomach. Similarly, mice were sacrificed via cervical dislocation and xenograft tumors collected. Tissues were fixed in 10% formalin, paraffin-embedded, and sections at 4μm interval were cut from each tissue, and stained with H&E, trichrome (Sigma Aldrich, St. Louis, MO), or via immunohistochemistry (IHC)/immunofluorescence (IF).

For IHC, slides were deparaffinized and heated in a pressure cooker using DAKO retrieval buffer (DAKO, Carpinteria, CA). Endogenous peroxidases were quenched in DAKO peroxidase block for 20 min. Tissues were blocked with 0.5% BSA in PBS for 30 min and exposed a primary antibodies against RAS^G12D^ (NewEast Biosciences, King of Prussia, PA), PCNA (Santa Cruz, Santa Cruz, CA), αSMA, Synaptophysin, Pancreatic amylase (abcam, Cambridge, MA), Vimentin, E-Cadherin, mutant P53, and pERK (Cell Signaling, Danvers, MA) at 1:50-1:400 overnight at 4°C. Slides were developed using an HRP-conjugated secondary antibody followed by DAB substrate/buffer (DAKO).

For IF, slides were deparaffinized and heated in a pressure cooker using DAKO retrieval buffer (DAKO, Carpinteria, CA). Endogenous peroxidases were quenched in DAKO peroxidase block for 20 min. Tissues were blocked with 0.5% BSA in PBS for 30 min and exposed a primary antibodies against CK19 (University of Iowa Hybridoma Bank, Iowa City, IA), pPancreatic amylase, αSMA, PCNA, (Santa Cruz), Vimentin, or pERK (Cell Signaling) at 1:100 overnight at 4°C. Slides were visualized using an Alexaflour-488 or 594 conjugated secondary (abcam).

For cultured cells, cells were grown on chamber slides and fixed with ice-cold methanol at −20^°^C for 10 minutes. Cells were blocked for 1 hour at room temperature with 0.5% BSA in PBS, and incubated with primary antibodies against CK19 (University of Iowa Hybridoma Bank), E-Cadherin, pERK, (Cell Signaling) or PCNA (Santa Cruz) at 1:50-100 overnight at 4°C. Slides were visualized using an Alexaflour-488 or 594 conjugated secondary antibody (abcam).

### Western blot and immunoprecipitation/RBD Assay

Cell or tissue lysates were lysed in 4% SDS buffer followed by needle homogenization. Equal amounts of protein (15–40 μg) were mixed with loading dye, boiled for 8 min, separated on a denaturing SDS–PAGE gel and transferred to a PVDF membrane. The membrane was blocked in 5% milk/TBS/0.1% Tween for 1 h and incubated with antibodies against pERK, ERK, Vimentin, (Cell Signaling, Danvers, MA, USA), αSMA (abcam), β-Actin (Santa Cruz Biotech, Santa Cruz, CA, USA). The membrane was washed with TBS-0.1% Tween and then incubated with HRP conjugated secondary antibody (Santa Cruz Biotech) at room temperature for 1 h and rewashed. Protein bands were visualized by an enhanced chemiluminiscence method (Thermo, Waltham, MA, USA) and resolved digitally per the manufacturer’s specifications. For RAS activation assay, a standard kit was purchased and used per manufacturer specification (Thermo). All experiments were performed in triplicate unless otherwise specified.

### Flow cytometry

Cultured pancreas cells were seeded into a round-bottom 96-well plate, washed in PBS, and stained with anti-RAS^G12D^ antibody (New East Biosciences) 1:100 in PBS with 1% BSA over ice for 20 minutes. Cells were analyzed with a BD Fortessa. All flow plots correspond to size appropriate single cells and are representative of 100,000 events unless otherwise stated.

### Study approval

All experiments involving the use of mice were performed following protocols approved by the Institutional Animal Care and Use Committee at the University of Illinois. Patient slides and information was obtained in a de-identified fashion from the Northwestern University Pathcore, following local IRB approval. All patients offered informed consent for study participation. All methods were performed in accordance with the relevant guidelines and regulations

## Results

### KRAS and TP53 mutations induce metastatic behavior in porcine pancreatic duct cells

To assess the viability of *loxp*-STOP-*loxp* (LSL)-KRAS^G12D^-TP53^R167H^ pigs as a potential model of PDAC, we first compared the microscopic anatomy of the human, murine, and porcine pancreas. Despite the described structural differences, we found that the gland histology was remarkably similar among the three species (Figure S1). In light of these similarities, we next euthanized a 2-month-old female LSL-KRAS^G12D^-TP53^R167H^ pig and removed the pancreas gland. We then extracted and digested the main pancreatic duct to isolate duct cells, and infected them with 1×10^6^ units of Adeno-Cre *in vitro.* Cells displayed normal ductal morphology and negligible proliferation 24 hours after transduction. However, 7 days following transduction, LSL-KRAS^G12D^-TP53^R167H^ expressing duct cells had self-immortalized, now displaying spindle shaped morphology consistent with malignant transformation (Figure 1A). Additionally, transduced cells also demonstrated robust expression of the TP53^R167H^ transcript/protein (Figure 1B,C and S2A), as well as the mutant RAS^G12D^ protein as determined by immunocytochemistry and verified by both western blot and flow cytometry (Figure 1D,E and S2A). These transformed cells, named Porcine Carcinoma 1 (PORC1), strongly expressed E-cadherin and CK19, affirming their ductal lineage. Consistent with a tumorigenic phenotype, PORC1 cells also displayed increased RAS activation, as well as strong expression of the KRAS effector pERK1/2 and proliferation surrogate PCNA (Figure 1F and S2B).

**Figure 1.**
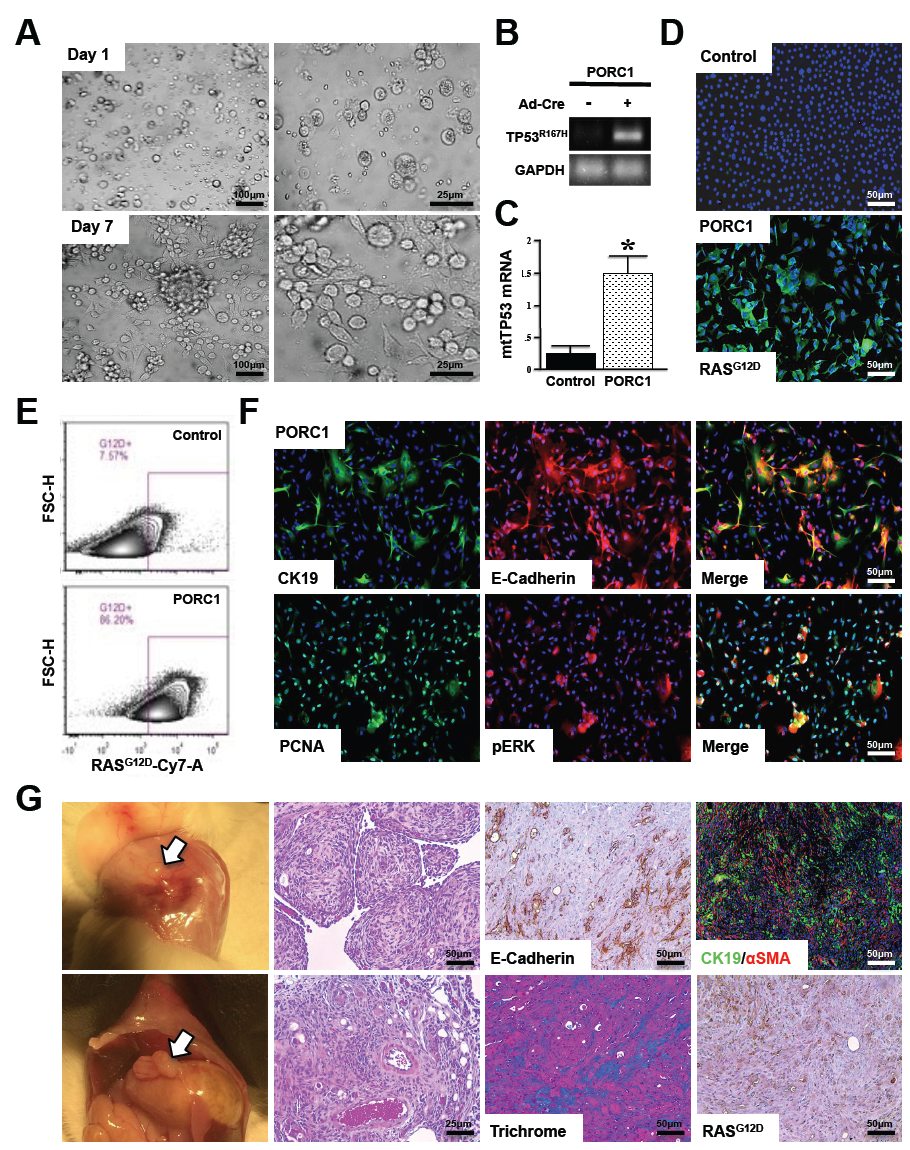
**Concomitant** ***KRAS^G12D^*** **and** ***TP53^R167H^*** **Mutations Induce Metastatic behavior in Porcine Pancreatic Duct Cells** ***ex vivo*** **(A)** Primary pancreatic duct cells were isolated, placed into culture, and incubated with 1×10^6^ units of Adenoviral-Cre. After 7 days the cells, designated Porcine Carinoma 1 or PORC1, self-immortalized and displayed more spindle shaped morphology **(B,C)** PORC1 cells were evaluated by RT-PCR for the TP53^R167H^ transcript. **(D,E)** Mutant RAS^G12D^ expression in PORC1 cells was confirmed using immunocytochemistry and flow cytometry. **(F)** PORC1 cells were validated using immunocytochemistry for the duct marker CK19, epithelial marker E-cadherin, as well as proliferation surrogate PCNA and the RAS effector pERK. **(G)** 5×10^6^ PORC1 cells were injected intraperitoneally into severe combined immunodeficient (SCID) mice. After 16 days, the mice presented with gross tumors at the site of injection, which were sectioned and stained with H&E, Masson’s Trichrome, or via immunohistochemistry for E-cadherin, CK19/αSMA, and RAS^G12D^ (N=7).

We next injected 5-10×10^6^ PORC1 cells either subcutaneously (SQ) or intraperitoneally (IP) into SCID mice (N=7 per group). Mice with SQ injection developed large masses under the skin at each injection site (Figure S3A), while mice with IP injection developed several large focal plaques on the abdominal viscera that were easily palpable (Figure 1G). SQ tumors were routinely measured and overall health was assessed by weight (Figure S3B). Like human PDAC, these tumors had a dense, cellular stroma, with cancer cells staining for E-cadherin, indicating an epithelial origin. These epithelial cells were distinct from the fibrous component, which was evaluated by Masson’s Trichrome staining. Furthermore, like the PORC1 cells *in vitro*, xenograft tumors were highly positive for CK19, affirming a ductal lineage and expressed mutant RAS^G12D^. Also consistent human PDAC tumors, the stroma had strong expression of αSMA, a marker of pancreatic stellate cells (Figure 1G and Figure S3A).

### Restriction of Adeno-Cre to the Main Pancreatic Duct Leads to a Predominantly Pancreatic Ductal Adenocarcinoma Histotype

Given the sufficiency of KRAS^G12D^ and TP53R167H mutations to transform pancreatic duct cells *in vitro*, we initiated autochthonous porcine pancreatic tumor formation *in vivo*. Our first attempts of intraparenchymal delivery of Adeno-Cre resulted in a mixed histotype of leiomyosarcoma and duct-derived neoplasms akin to those seen in human patients (N=3, Figure S4 and S5, and Supplementary Results). Given the presence of clinically relevant pancreatic neoplasms, we next attempted to recreate the described duct-derived pancreas histotype pancreas histotype while avoiding the leiomyosarcoma phenotype observed in our initial experiments. Therefore, we repeated the above experiment focusing on transformation of ductal epithelial cells rather than mesenchyme by surgically injecting 4×10^9^ units of Adeno-Cre directly into the main pancreatic duct.

One year following injection, there were no overt signs of illness or pancreatic insufficiency as observed in the previous cohort. Similarly, contrast-enhanced computed tomography (CT) showed no evidence of gross tumor formation at the injection site or near the pancreas. Additionally, the hepatobilliary tract, spleen, liver, and regional lymph nodes appeared normal with no sign of lesions. The stomach was moderately filled with gas and fluid with a mild amount of hyper-attenuating material in the dependent region of the fundus, though there was no clear sign of overt abnormality (Figure S6A). Furthermore, upon euthanasia, the pancreas displayed no overt signs of sarcoma and had three distinct lobes. However, on dissection, the main pancreatic duct was large and firm with several large, nodular tumors with a pronounced fibrous component at the site of cannulation (Figure S6B).

Tumors were then fixed and sectioned, and compared directly to human PDAC samples. Upon histological evaluation of several geographically separate areas, porcine tumors consistently displayed morphological features consistent with human PDAC (Figure 2A,B). In addition to having no evidence of sarcoma or smooth muscle abnormality, there were several areas consisting of lumenal masses with considerable cellular atypia and invasion through the basement membrane (Figure 2B). Like human PDAC, these lesions were accompanied by a dense and desmoplastic tumor stroma rich with leukocytes (Figure 2A,B).

**Figure 2.**
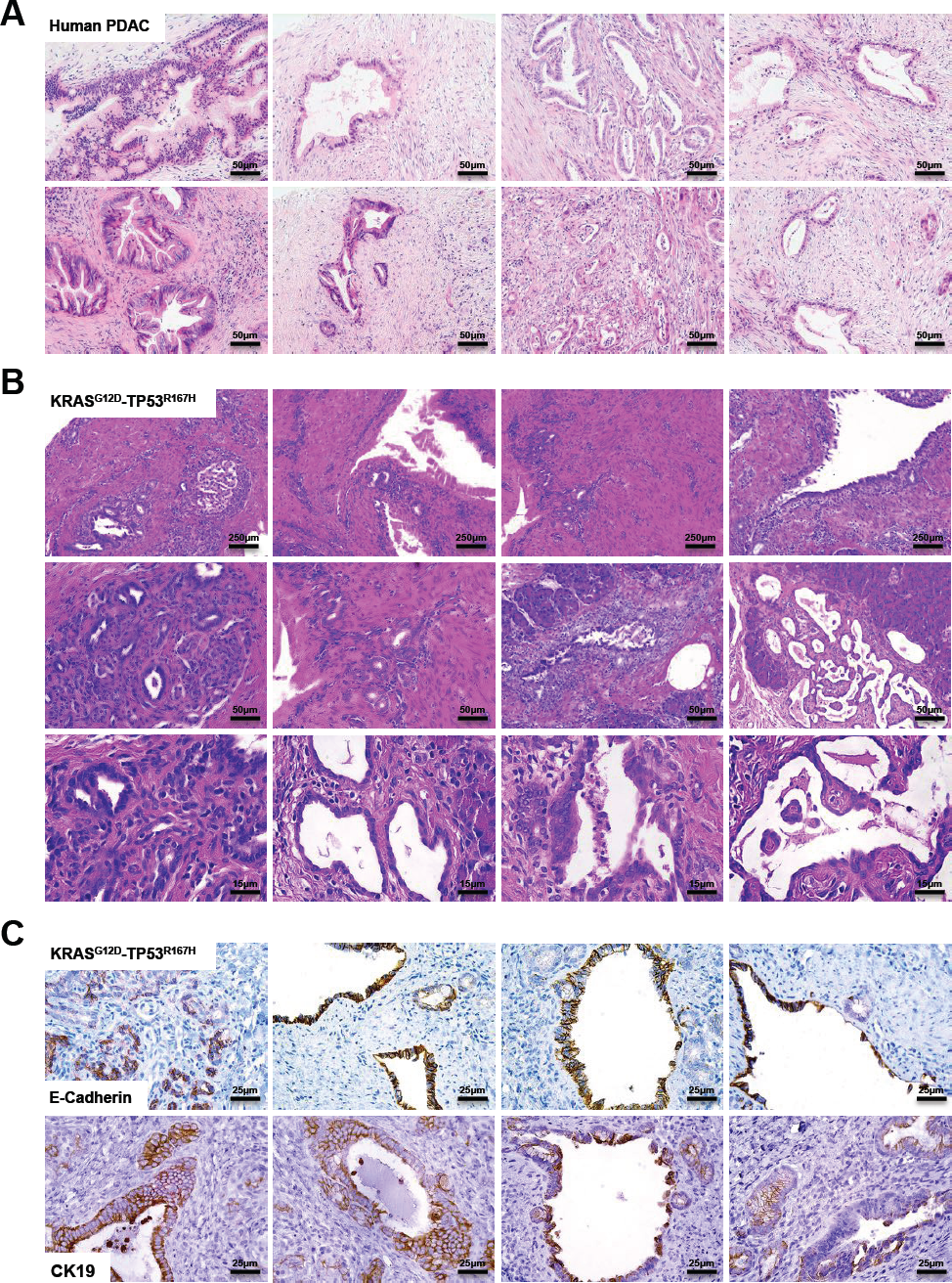
**Restriction of Adeno-Cre to the Main Pancreatic Duct Leads to a Predominantly Pancreatic Ductal Adenocarcinoma Histoype (A)** Sections from human PDAC patients were stained with H&E demonstrating varied tissue architecture with common features including ductal lesions with cellular atypia and a dense tumor stroma. **(B)** Tumors from the LSL-KRAS G12D-TP53^R167H^ pig delivered an Adeno-Cre injection into the main pancreatic duct were sectioned and stained with H&E showing several histologic features consistent with PDAC. **(C)** Porcine tumors were next stained for the pan-epithelial marker E-cadherin or the duct marker CK19, affirming a PDAC histotype.

Consistent with a PDAC phenotype, porcine tumor sections were positive for both E-cadherin and CK19 (Figure 2C). Though the predominant histotype was derived from exocrine pancreas, there were distinct and geographically separate areas that appeared to have a phenotype more similar to pancreatic neuroendocrine tumors (PNET). We therefore compared these areas directly to human PNET sections, and found that they shared several histological features, including a less pronounced desmoplastic tumor stroma (Figure S7A,B) and stained positive for the neuroendocrine marker Synaptophysin (Figure S7C).

### LSL-KRAS^G12D^-TP53^R167H^-Induced PDAC Displays Excessive Proliferation

To assess changes in KRAS activity, we first performed a RAS activation assay using control and tumor tissues. As expected, in PDAC tissues KRAS protein had increased GTP binding ratio compared to control tissues, and exhibited expression of KRAS^G12D^ and P53^R172H^ proteins. These were accompanied by enhanced ERK activation as determined by western blotting (Figure S8) and confirmed by immunohistochemistry, localizing pERK to areas suspect for PDAC (Figure 3A and S9A). These areas also stained strongly for PCNA, a surrogate marker of cell proliferation (Figure 3B and S9B), whereas adjacent normal tissue had little to no PCNA staining (data not shown). To affirm that these proliferating regions were indeed those previously identified as PDAC, we dual-stained tissues for both the ductal marker CK19 and PCNA. Regions proliferating most strongly were also CK19 positive, indicative of PDAC (Figure 3C). Similarly, areas suspect for PNET also displayed robust ERK activation as well as PCNA staining, further suggesting the presence of endocrine-derived neoplasms (Figure S7D).

**Figure 3.**
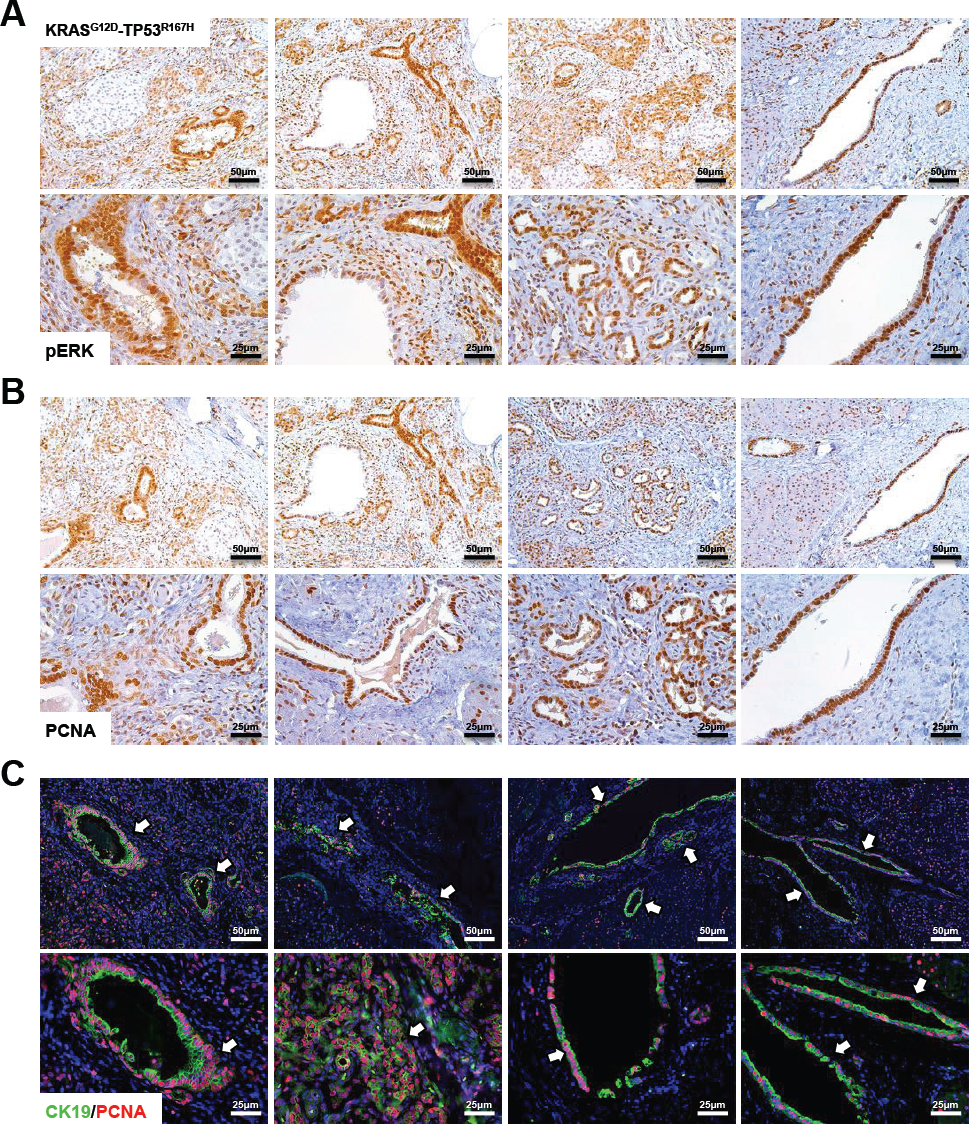
**LSL-KRAS^G12D^-TP53^R167H^-Induced PDAC Displays Excessive Proliferation (A,B)** To assess downstream RAS activity and cell proliferation, porcine tissues were stained via immunohistochemistry for pERK or PCNA, and compared to non-infected pancreas tissue of LSL-KRAS^G12D^-TP53^R167H^ pigs. Sections stained for pERK were scored based on intensity and scores averaged, showing increased ERK phosphorylation. Similarly, PCNA^+^ cells were quantified per 40X and compared to non-infected controls, indicating an increase in cell proliferation. **(C)** Sections were dual stained for CK19 and PCNA to localize cell proliferation to areas of PDAC. Dual positive cells per 40X field were then quantified and compared to non-infected pancreas tissue of LSL-KRAS^G12D^-TP53^R167H^ pigs.

### Porcine PDAC Develops a Desmoplastic Stroma Analogous to Human Patients

As the pancreatic cancer stroma is not only a near uniform histological feature but also a key component of disease progression and chemo-resistance^17,18^, we analyzed the tumor microenvironment of both human and porcine PDAC sections. As expected, all human PDAC samples examined had a dense, desmoplastic tumor stroma that stained positive with Masson’s Trichrome (Figure 4A). Porcine tumors were similarly desmoplastic with increased Collagen IA expression (Figure S8) and also stained positive with Masson’s Trichrome (Figure 4B). Similarly, expression of the mesenchymal marker Vimentin was increased (Figure S8) and localized exclusively to the tumor stroma (Figure 4C). As pancreatic stellate cells are considered the critical mediator of tumor-associated fibrosis^19^, we next assessed tumor sections for expression of αSMA, which was also localized to the stroma and was associated with areas of CK19-positive PDAC (Figure 4D).

**Figure 4.**
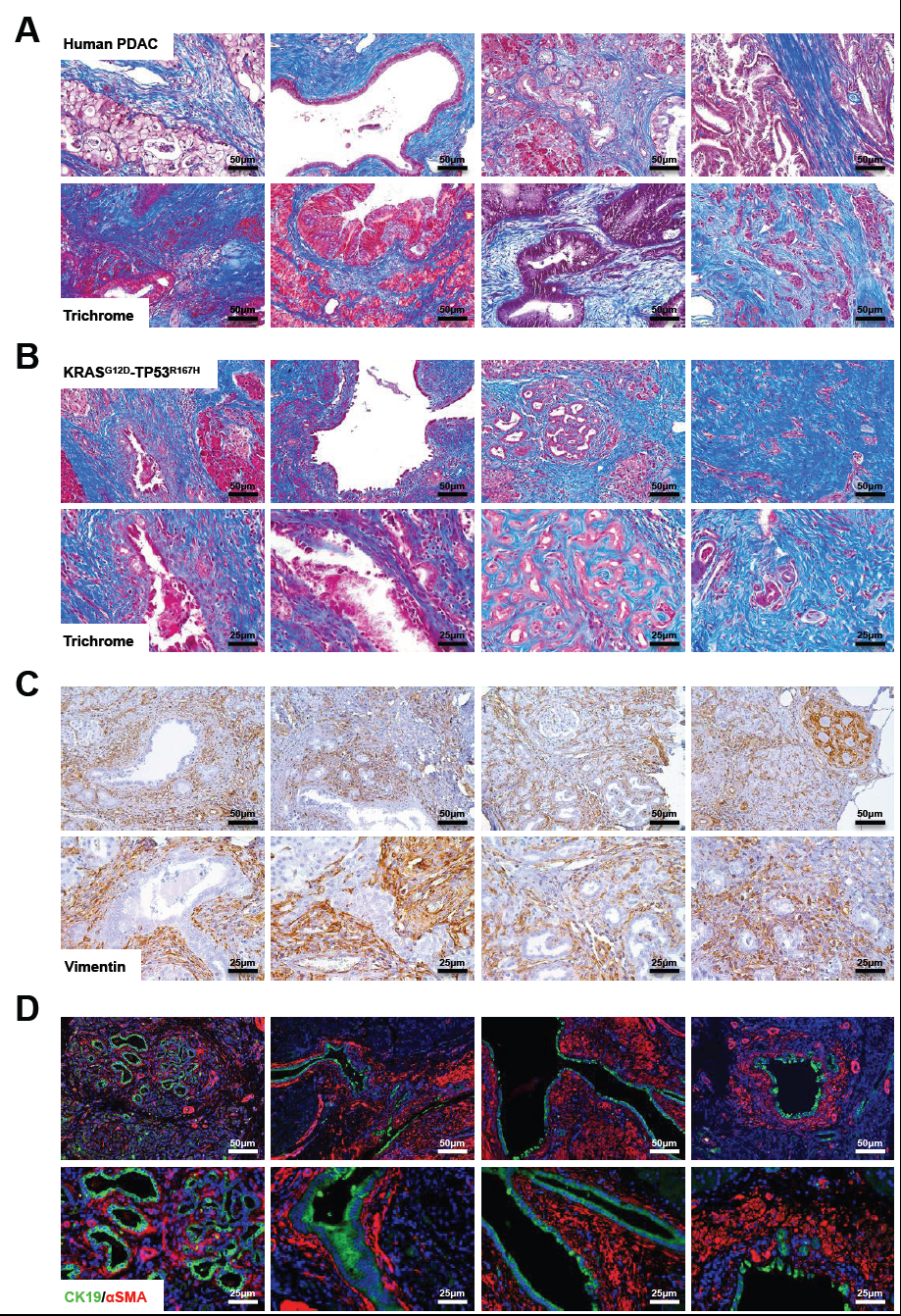
**Porcine PDAC Develops a Desmoplastic Stroma Analogous to Human Patients (A,B)** The tumor microenvironment from human PDAC patients and duct-injected LSLKRAS^G12D^-TP53^R167H^ pigs were assessed via Masson’s Trichrome staining. **(C)** Pig tissue sections were stained for the mesenchymal marker Vimentin. **(D)** Sections were dual stained for CK19/αSMA to evaluate the presence of pancreatic stellate cells surrounding PDAC lesions.

## Discussion

The poor survival associated with pancreatic cancer stems largely from the fact that most patients initially present with highly advanced disease. As most of these patients are not candidates for surgical intervention and there is no effective pharmacological therapy, there is an urgent need to develop new therapeutic approaches. To do so, investigators have relied almost exclusively on genetically engineered mouse models. For example, the Pdx promoter has been used to drive Cre expression in the progenitor cells of the pancreas, which when crossed to a *loxp*-STOP-*loxp* (LSL) cassette resulted in a faithful recapitulation of the more common PanIN histotype^20^. Combining this model with an LSL-TP53^R172H^ cassette resulted in a metastatic model of PDAC with histology closely resembling that observed in human patients^10^. These KPC mice have been favored by a number of investigators for this reason, and have provided a reliable and accurate model of human PDAC for over a decade. However, despite the success of the KPC mice, there are still the inherent limitations of the mouse anatomy and physiology that must be considered.

Importantly, the mouse pancreas has significant micro and macroscopic differences to that of humans. The human pancreas is a firm organ that is segmented into three borderless, yet distinct parts: the head, body, and tail^21^. In contrast, the mouse pancreas is soft and diffuse with three poorly defined lobes^22^. There are also pronounced differences of the ductal tree between the two species. In humans, the pancreatic acini are the predominant exocrine cell type and are organized into lobules that secrete to a small, intercalated duct. These intercalated ducts drain to a larger, interlobular duct that join to form the large, main pancreatic duct which empties to the duodenum adjacent to the common bile duct^21^. In mice, however, a large interlobular duct drains the three respective lobes. Ducts from the splenic and gastric lobes then merge with the common bile duct far more proximal to the duodenum than in humans^21^.

In addition to these structural variances, there are several genetic factors that account for further differences in mouse and human carcinogenesis. Despite sharing a similar number of protein coding genes^23^, the mouse genome is substantially different from that of humans. Though much is shared, there is significant variance in transcriptional regulation and chromatin organization between the two species^24^. This is particularly true of genes involved in the inflammatory response^25^, indicating that mice may have limited utility in inflammation-associated diseases such as pancreatic cancer^26^. Another notable disparity between mouse and human genetics is the failure of mutations in Cystic Fibrosis Transmembrane Regulator (CFTR) gene to induce significant pancreatic disease as it does in humans^27^. These issues were resolved by modeling the disease in pigs, which lead to a more faithful recapitulation of the disease pathology and destruction of the exocrine pancreas^28,29^. We therefore attempted to recreate the success of the KPC model in the more genetically and anatomically similar *sus scrofa* domestic pig, in order to provide a more clinically relevant model of PDAC.

Initial attempts at intrapancreatic delivery of Adeno-Cre resulted in a mixed histotype. While the pigs developed PanIN lesions within 16 days, we also observed lethal smooth muscle derived leiomyosarcoma, likely due to the malignant transformation of vascular smooth muscle infected with Adeno-Cre. These two cancer types generally do not present simultaneously. Therefore, it was necessary to refine our approach in order to generate a clinically relevant PDAC histotype. By restricting the delivery of our Cre using an intraductal injection, we were able to induce locally invasive PDAC without the presence of an underlying sarcoma. Interestingly, this approach also resulted in a somewhat mixed histotype, this time consisting of PDAC as well as rare and geographically isolated regions of PNET.

Mixed tumors and mixed adenoneuroendocine carcinomas (MANEC) of the gastrointestinal tract and pancreas have been well described by our group as well as many others^30^. They are commonly aggressive tumors most frequently occurring in the head of the gland and management is often a combination of surgery and platinum-based chemotherapy. At the present time, this may be the most appropriate category for tumors induced by LSLKRAS^G12D^-TP53^R167H^ pigs, though PDAC appeared to predominate, particularly in sections closest to the main pancreatic duct. Therefore, future work will employ improved exocrine targeting to exclude the neuroendocrine compartment.

Based on the size and anatomy of the pig, this model may allow for insight into surgical and interventional radiology techniques never before possible in rodents. Pigs and humans also have largely analogous Cytochrome P450 enzymes, allowing for better pre-clinical evaluation of drug metabolism and therapy^31^. Additionally, given the profound similarity between human and pig immune systems, these LSL-KRAS^G12D^-TP53^R167H^ pigs may provide an improved platform for preclinical immunotherapy. Given these advantages, several porcine models of cancer are emerging. These include a mutant APC porcine model of familial adenomatous polyposis (FAP), a heterozygous TP53 knockout model of spontaneous osteosarcomas, and a chemically induced hepatocellular carcinoma (HCC) model^32^. Similarly, our inducible LSL-KRAS^G12D^-TP53^R167H^ model has successfully generated soft tissue sarcomas and HCC^16,32,33^.

By demonstrating sufficiency of the LSL-KRAS^G12D^-TP53^R167H^ pig to model a human-like pancreatic carcinoma both *in vitro* and *in vivo* (Summarized in Figure 5), we substantiate the pig as a more physiologically relevant platform in which to model the two predominant pancreatic cancer histotypes, allowing for novel research approaches and bridging the gap from bench to bedside.

**Figure 5.**
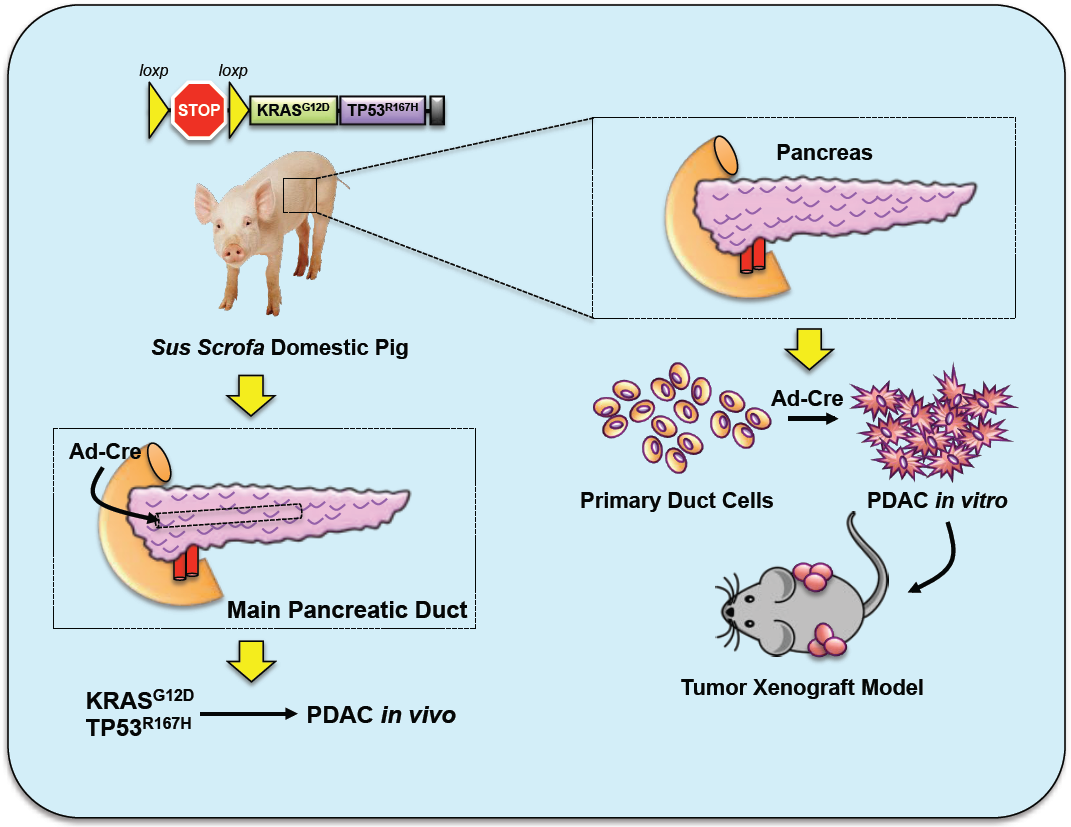
**Schema Summarizing the LSL-KRAS^G12D^-TP53^R167H^ porcine model of PDAC** By isolating pancreatic duct cells from LSL-KRAS^G12D^-TP53^R167H^ pigs and transducing with Adeno-Cre *in vitro*, cells self-immortalized and gained the ability to establish tumors in xenograft experiments using immunocompromised mice. Similarly, when Adeno-Cre was administered to the main pancreatic duct *in vivo*, LSL-KRAS^G12D^-TP53^R167H^ pigs developed distinct PDAC along the injection site.

## Acknowledgments

This work is dedicated to the memory of our friend and mentor Dr. Howard Ozer MD, PhD.

## Abbreviations

Adeno-Cre: Adenoviral Cre-Recombinase
PanIN: Pancreatic Intraepithelial Neoplasm
NET: Neuroendocrine Tumor
PDAC: Pancreatic Ductal Adenocarcinoma

